# Tissue osmotic surveillance shapes susceptibility of zebrafish to wound infections

**DOI:** 10.64898/2026.08.04.742764

**Authors:** Ivanna Williantarra, Salik Miskat Borbora, Hazel A. Walker, Auxence Desrentes, Milka Sarris

## Abstract

Wounds represent compromised tissues that are susceptible to opportunistic infections. Tissue factors predisposing to wound infection include edema, poor vascularity, tissue hypoxia, necrosis, and wound size. However, it is unclear whether factors external to the tissue play a role. Detection of osmolarity imbalances between internal and external environment has been shown to alter epithelial wound closure and immune cell recruitment at injury sites. However, it remains unclear whether these osmotic surveillance pathways affect susceptibility to pathogenic wound infections. Here, we use live imaging and disease monitoring in zebrafish larvae to understand susceptibility of wound infection towards *Pseudomonas aeruginosa*. We show that exposure of wounds to isotonic solution results in enhanced susceptibility and pathogen burden. Furthermore, we demonstrate that this higher propensity for infection depends on osmolarity-mediated changes in wound sealing and neutrophil recruitment during the early stages of the wound response. These findings inform the design of experimental wound infection models as well as the clinical management of wounds.

## Introduction

Wounds occur when tissue integrity is compromised upon disruption of anatomical architecture with concomitant loss of normal physiological function [1]. They require efficient resolution as inadequate repair mechanisms could lead to serious complications such as persistent inflammation, pathogen growth, tissue necrosis and sepsis. Wound infections remain a major healthcare challenge in acute and chronic wounds, contributing significantly to patient morbidity, prolonged hospitalization, and mortality. As per World Health Organization estimates, 11% of patients undergoing surgery in low- and middle-income countries develop a surgical site infection (SSI) (www.who.int). Additionally, chronic wound cases such as diabetic foot ulcers, venous leg ulcers, and pressure injuries are prone to pathogenic infection [2], substantially impairing health and life. Thus, the global wound management expenditure has significantly increased in the last decade, reaching a staggering $148.65 billion in 2022 [3].

*Pseudomonas aeruginosa* is an opportunistic bacterial pathogen, ubiquitously present in nature, able to colonise wounds in multiple species. In humans, wound infections can occur through contact of injured tissue with contaminated water or surfaces such as contaminated hot baths or spa pools. These infections can be severe and lead to sepsis and mortality, particularly in immunocompromised individuals or in the case of large surgery-associated infections, [4, 5]. *P. aeruginosa* is one of the top-listed pathogens causing hospital-acquired infections owing to their reported presence on medical devices as they thrive on wet surfaces [6]. A host of virulence factors contribute towards a successful *P. aeruginosa* infection. This includes lipopolysaccharide (LPS), responsible for antibiotic tolerance [7], out-membrane proteins contributing to adhesion and nutrient exchange [8], secretion systems involved in colonization of the host [9], and exopolysaccharides, involved in impairment of bacterial clearance [10].

Host tissue responses also influence the scale of *Pseudomonas* infections.Wounded regions require efficient sealing to restore barrier integrity, prevent fluid loss, protect against infection, and enable regeneration and tissue repair [11, 12]. Following tissue injury, epithelial cells at the site rapidly detect damage and become activated. They move across the wound matrix to envelop the breached surface [13, 14]. This is followed by active cell division to ensure sufficient cell numbers for wound closure. Once the wound is sealed, keratinocytes differentiate into organized epidermal layers, restoring normal tissue architecture and function [15, 16]. Additionally, epithelial cells orchestrate the early recruitment of immune cells, the earliest of which are neutrophils [15]. Epithelial barrier disruption produces changes in tissue tension and mechanical stress, leading to activation of intracellular signaling cascades and production of inflammatory mediators that promote neutrophil recruitment [17]. Calcium signaling propagating from damaged cells into the surrounding tissue stimulates chemoattractant release that aids neutrophil recruitment [18]. Furthermore, upon tissue barrier breach, damage-associated molecular patterns, such as adenosine triphosphate (ATP), nucleic acids, formylated peptides further add to the chemoattractant cocktail for neutrophil recruitment [19].

The influence of the external environment on wound infection susceptibility is less clear. Osmotic surveillance has been shown to play a role as a damage detection pathway that influences wound sealing and immune cell recruitment in sterile wounds [20]. However, how this pathway influences wound infection susceptibility is unclear. Previous studies suggest that tissue damage detection by osmotic surveillance supports responses to micro-injected *Pseudomonas aeruginosa*[21]. However, since micro-injection of bacteria bypasses the natural colonisation route, it remains unclear how much this damage detection pathway contributes to the progression of wound infections. Here, we address this by exploiting the transparency and tractability of zebrafish that allows visualization of innate immune cells in real time[22, 23]. Additionally, zebrafish larvae has also been employed to model wounds in specific conditions [24]. Using two distinct modes of tissue injury, we report that exposure to isotonic buffer conditions accompanied robust pathogen colonization and worsened disease course. Furthermore, osmotic differences contributed to disparity in neutrophil recruitment at the wound sites and altered wound closure dynamics in zebrafish larvae.

## Results

### Detection of osmotic imbalance limits susceptibility to wound infection *by Pseudomonas aeruginosa*

To investigate the role of osmotic surveillance in wound infections, we employed a previously introduced model wherein zebrafish larvae are subjected to wounding in the presence of aqueous medium contaminated with *P. aeruginosa* PAO1 strain [18]. This experimental model is advantageous in that it recapitulates a natural route of waterborne infection, in comparison to more common microinjection routes [21], and provides a relevant setting to study the role of immune cell dynamics in the initial stages of the wound in driving infection susceptibility.

To test the role of osmotic surveillance, we subjected zebrafish larvae to mechanical wounding (**Fig. 1A**), in two common zebrafish buffers E3 medium (hypotonic), Ringer medium (isotonic), as well as isotonic E3 medium (with the same sodium levels and tonicity as Ringer). We found that larvae in hypotonic medium were largely resistant to wound infection, whereas larvae in isotonic medium (iso E3, Ringer) were susceptible (**Fig. 1C**, *left panel*). In all media conditions, non-wounded larvae were resistant to infection, confirming the wound-dependent nature of infection (**Fig. 1C**, *right panel*). Larvae infected in hypotonic media showed minimal bacterial burdens in comparison to larvae infected in isotonic media (**Fig 1D**). To understand disease susceptibility in a granular manner, we characterised development of disease in individual larvae by creating a disease scoring matrix. Each larva was scored *(From 0-4, with a higher number indicating aggravated disease pathology*) based on phenotypic parameters that ranged from alterations in swimming behaviour, to more extensive morphological abnormalities viz. degree of tissue disintegration and cessation of heartbeat **(Fig. 1B)**. Infection in hypotonic media resulted in minimal development of disease across the 24 h period of observation (**Fig. 1E**, *left panel*). In contrast, exposure to PAO1 under isotonic conditions induced a pronounced disease phenotype in larvae over the entire 24-hour time course, with onset of disease evident at earlier timepoints of observations (**Fig. 1E**, *middle and right panel*).

**Figure 1.**
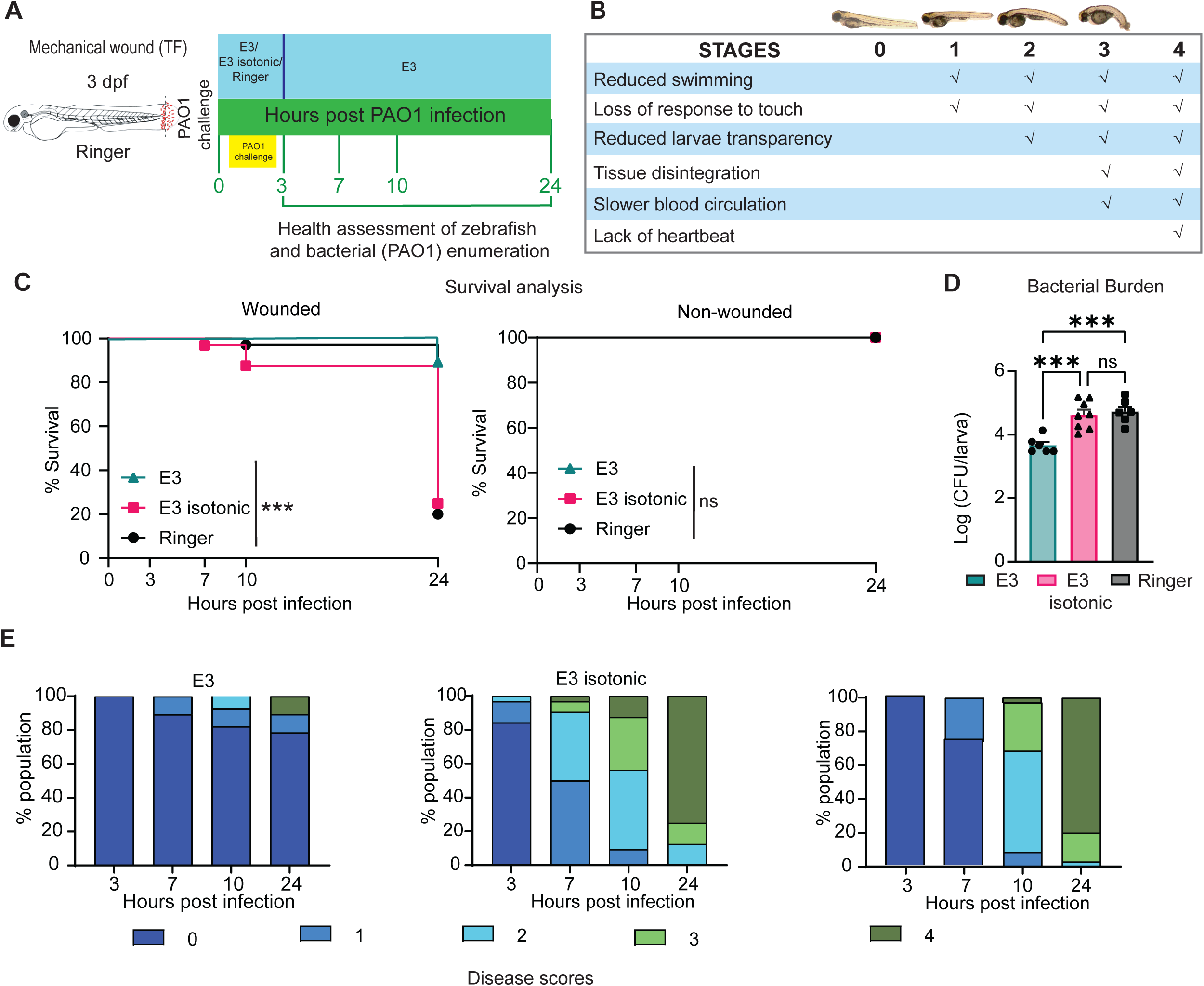
Osmotic surveillance determines susceptibility to mechanical wound infection. A. Schematic detailing the mechanical wound (MW) infection model by tail fin (TF) transection method in the presence of *P. aeruginosa* (PAO1). B. Parameters assessed in the disease scoring matrix to accord a disease progression score to each zebrafish larva. C. Survival curves from zebrafish larvae when exposed to indicated buffers and *P. aeruginosa* infection upon MW. (Representative of three independent experiments with n ≥ 24 larvae per group for the wounded-infected; n ≥ 9 larvae per group for the non-wounded infected, in each experiment). Log-rank (Mantel-Cox) test; ***P < 0.001 D. Bacterial burden of *P. aeruginosa* in the specified groups of zebrafish larvae at 24 hpi (hours post infection). CFU, colony forming units. (Representative of three independent experiments with n ≥9 larvae per group used for CFU analysis in each experiment). (mean ± SD). One-way ANOVA with Tukey’s multiple comparisons test, ***P < 0.001 E. Disease progression in the specified group of larvae at the indicated time points. Representative of three independent experiments, with n ≥ 24 larvae per group.

### Early tissue response to osmotic imbalance plays a role in susceptibility to wound infection

Wound closure can be delayed in isotonic media due to lack of tissue damage detection by the ruptured epithelium [25]. We hypothesised that this may render wounds more permissive to bacterial colonisation, contributing to the increased burden of infection in such conditions. To test whether early wound colonisation might vary in the different buffers, we measured bacterial burden shortly after wound infection (10 min and 30 min), at time points shorter than the duplication time of *P. aeruginosa* [26]. We found that at this early stage, the bacterial titres were already significantly higher in isotonic conditions **(Fig. 2A)**, suggesting a difference in tissue response in the initial stages of infection.

**Figure 2.**
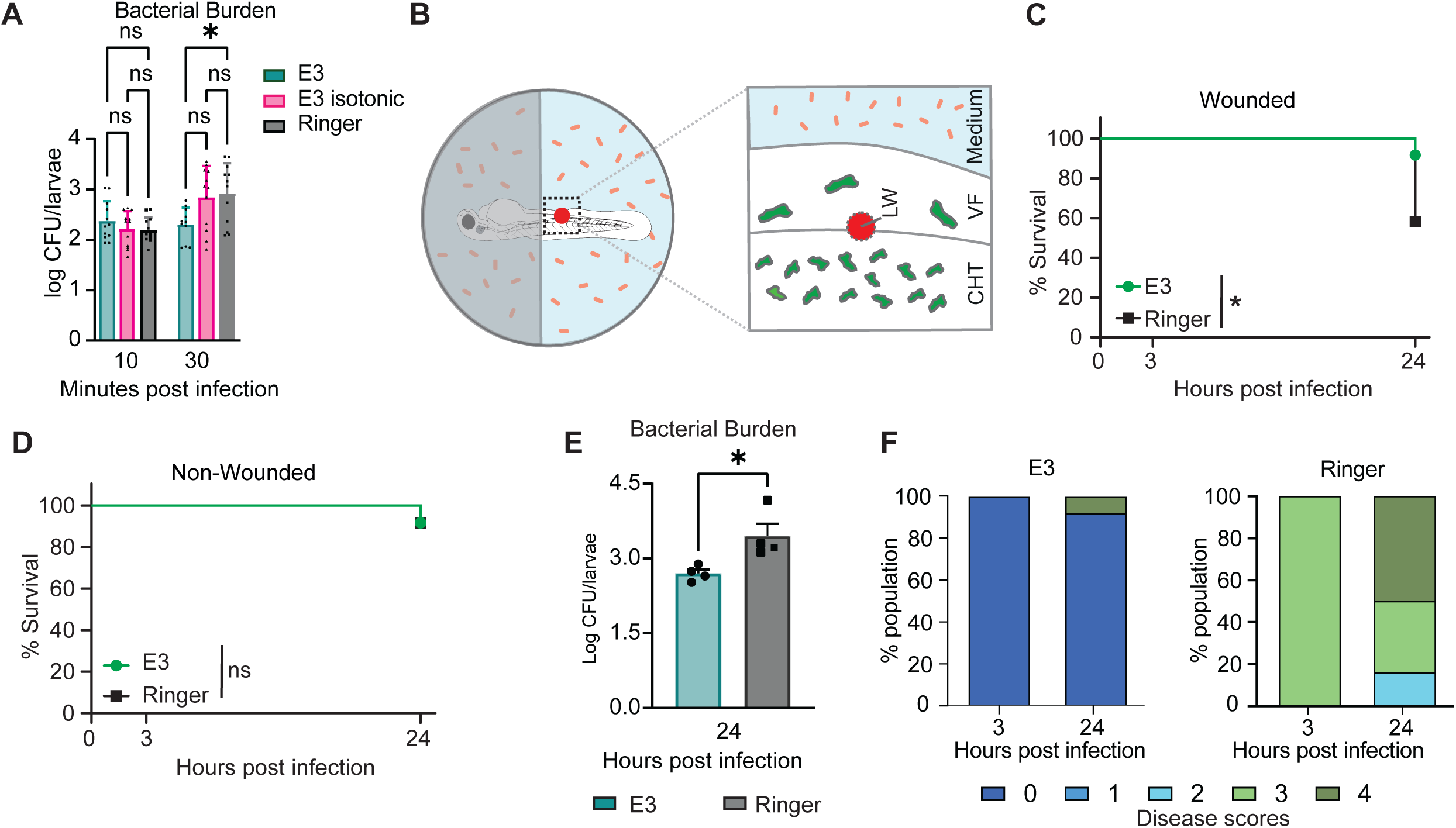
Osmotic surveillance determines susceptibility to laser wound infection. A. Bacterial burden of *P. aeruginosa* in the specified groups of zebrafish larvae at early time points (10 mins and 30 mins) of infection. (Representative of three independent experiments with n ≥ 12 larvae per group per time point in each experiment. (mean ± SD). CFU, colony forming units. One-way ANOVA with Sidak’s multiple comparisons test, **P < 0.01 B. Schematic detailing the laser wound (LW) infection model. CHT: caudal hematopoietic tissue; VF: ventral fin. C. Survival curves from zebrafish larvae when exposed to indicated buffers and *P. aeruginosa* infection upon LW. (Representative of three independent experiments with n ≥ 24 larvae per group in each experiment). Log-rank (Mantel-Cox) test, *P < 0.05 D. Survival curves from zebrafish larvae when exposed to indicated buffers and *P. aeruginosa* infection in the absence of LW. (Representative of three independent experiments with n ≥ 12 larvae per group in each experiment) Log-rank (Mantel-Cox) test, ns: non-significant E. Bacterial burden of *P. aeruginosa* in the specified groups of zebrafish larvae at 24 hpi (hours post infection). Representative of three independent experiments, with n ≥12 larvae per group in each experiment. (mean ± SD). CFU, colony forming units. Unpaired t test, *P < 0.05 F. Disease progression in the specified group of larvae at the indicated time points. Representative of three independent experiments with n ≥ 24 larvae per group.

To visualise the initial response to wound infection, we employed laser wound infection model, [18]. To extend our findings to this assay, we tested zebrafish susceptibility to PAO1 infection in hypotonic (E3) buffer and isotonic (Ringer) buffer respectively. Briefly, anesthetised larvae were mounted in low-melting-point agarose and immersed in a bacterial suspension prepared in distinct buffers. This allowed wound-associated infection to occur immediately after laser injury was inflicted (**Fig. 2B**). In line with the mechanical wound assay, infection in hypotonic media upon laser wounding elicited resistance in zebrafish larvae when compared to isotonic infection conditions (**Fig. 2C**). Non-wounded control larvae did not show any differences, indicating the requirement of wounds for infection establishment (**Fig. 2D**). Isotonic conditions resulted in robust bacterial burden (**Fig. 2E**) and exacerbated disease progression when compared with infection in hypotonic conditions, even from the early 3-hour time point (**Fig. 2F**). The commensurable infection outcome and bacterial burden in isotonic environment across the two wound models suggest that greater sensitivity to infection in those conditions arise from changes in early host immune responses, enabling potent bacterial colonisation.

### Osmotic imbalance detection limits bacterial colonisation

Microbial colonisation is limited by rapid wound closure. We reasoned that higher bacterial counts in the early time points of infection could be a result of disparate wound sealing in the different buffers. To confirm the previously described effects of tonicity on wound closure [25], we used the localised laser wound assay in sterile conditions. Wound closure (**Fig. 3A,B, Movie S1**) and neutrophil recruitment (**Fig. 3C**) was delayed in isotonic buffer under sterile solutions. Since osmolarity differences can also influence *Pseudomonas aeruginosa* physiology [27], we tested direct effects of the buffer compositions on the growth of bacteria. We found comparable bacterial growth after 30 min incubation in the said media (**Fig. 3D**), suggesting that the differential microbial colonisation in isotonic versus hypotonic solutions is not due to bacterial fitness differences. Together, these data indicated that either slower wound closure or sub-optimal recruitment of neutrophils towards injury sites in isotonic conditions drive disease.

**Figure 3.**
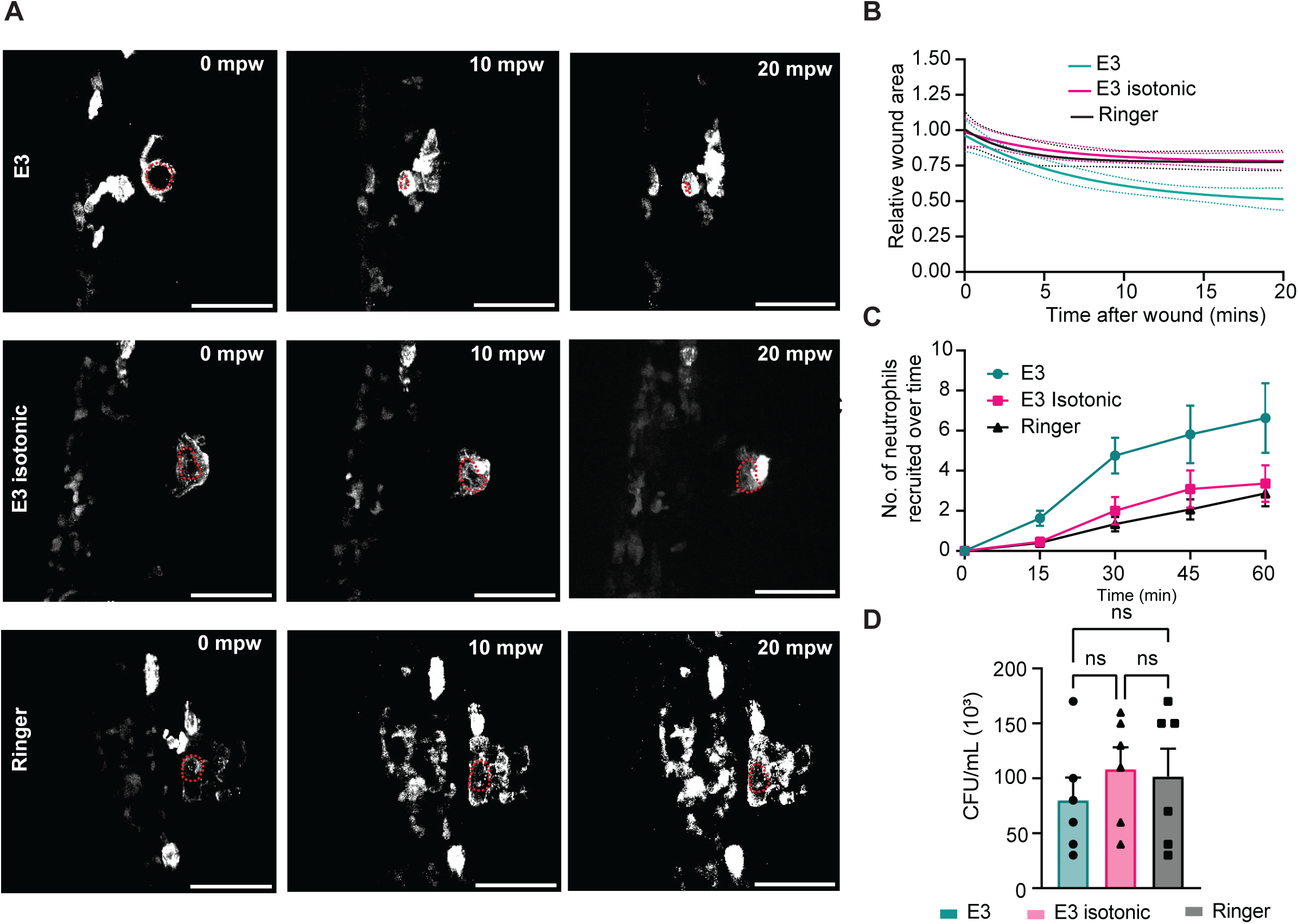
Differential osmolarity detection affects epithelial and immune responses during laser injury in zebrafish larvae. A. Time lapse images of representative movies of zebrafish larvae at the indicated time points relative to the time of LW. 0 mpw *(left panel)*, 10 mpw *(middle panel),* and 20 mpw *(right panel)*. Red dashed lines correspond to the wound margin. LW: laser wound, mpw: minutes post wounding. Scale bar 50 μm. B. Comparison of the changes in wound area across three buffers (E3, E3 isotonic and Ringer). Relative wound area was measured over time for laser wounds created in zebrafish larvae incubated in E3, E3 isotonic and Ringer buffer. Wound closure was monitored over time. Solid lines indicate non-linear regression fits using a one-phase exponential decay model. While individual goodness-of-fit was low (E3 : R squared=0.1044, SS=6.057; E3 isotonic: R squared=−0.076,SS=3.453; Ringer: R squared=0.02,SS=2.677), with a global cumulative R squared=0.1721, an Extra Sum-of-Squares F-test (Total SS = 12.19) indicate that the curves were different between at least one pair of conditions (P<0.0001). n =10 (E3), n=9 (E3 isotonic), n=7 (Ringer). SS: sum of squares. Scale bar: 50 μm C. Number of neutrophils recruited at the wound site in zebrafish larvae incubated in E3, E3 isotonic and Ringer buffer after laser wound (LW) in sterile conditions. n=16 (E3), n=11 (E3 isotonic), n=15 (Ringer). Two-way ANOVA with Tukey’s multiple comparisons test. No statistical differences in neutrophil numbers in comparisons across the three buffers. Within the same buffer, there was a difference between the number of neutrophils recruited at 60 min. over 15 min. D. Bacterial PAO1 (*P. aeruginosa*) counts after incubation of the microbe in the indicated buffers for 30 min. Representative of three independent experiments with six replicates per group in each experiment. CFU, colony forming units. (mean ± SD). One-way ANOVA with Tukey’s multiple comparisons test, ns: not significant

Unlike in sterile settings, neutrophil recruitment to infected wounds is stimulated by both damage detection and bacterial detection [19]. We reasoned that increased bacterial colonisation due to delayed wound sealing in isotonic conditions results in different neutrophil recruitment kinetics in comparison to sterile wounds. To test this, we quantified neutrophil recruitment in wounds exposed to PAO1 (**Fig. 4A, Movie S2**). In contrast to sterile settings, the magnitude of overall neutrophil recruitment was similar in hypotonic and isotonic conditions (**Fig. 4B**). This was a result of a delayed increase in neutrophil recruitment after the first 30min in isotonic conditions, which was not observed in hypotonic conditions (**Fig. 4C,D**). This delayed rise in neutrophil recruitment in infected wounds was consistent with higher bacterial detection after the first 30 min of wound-infection.

**Figure 4.**
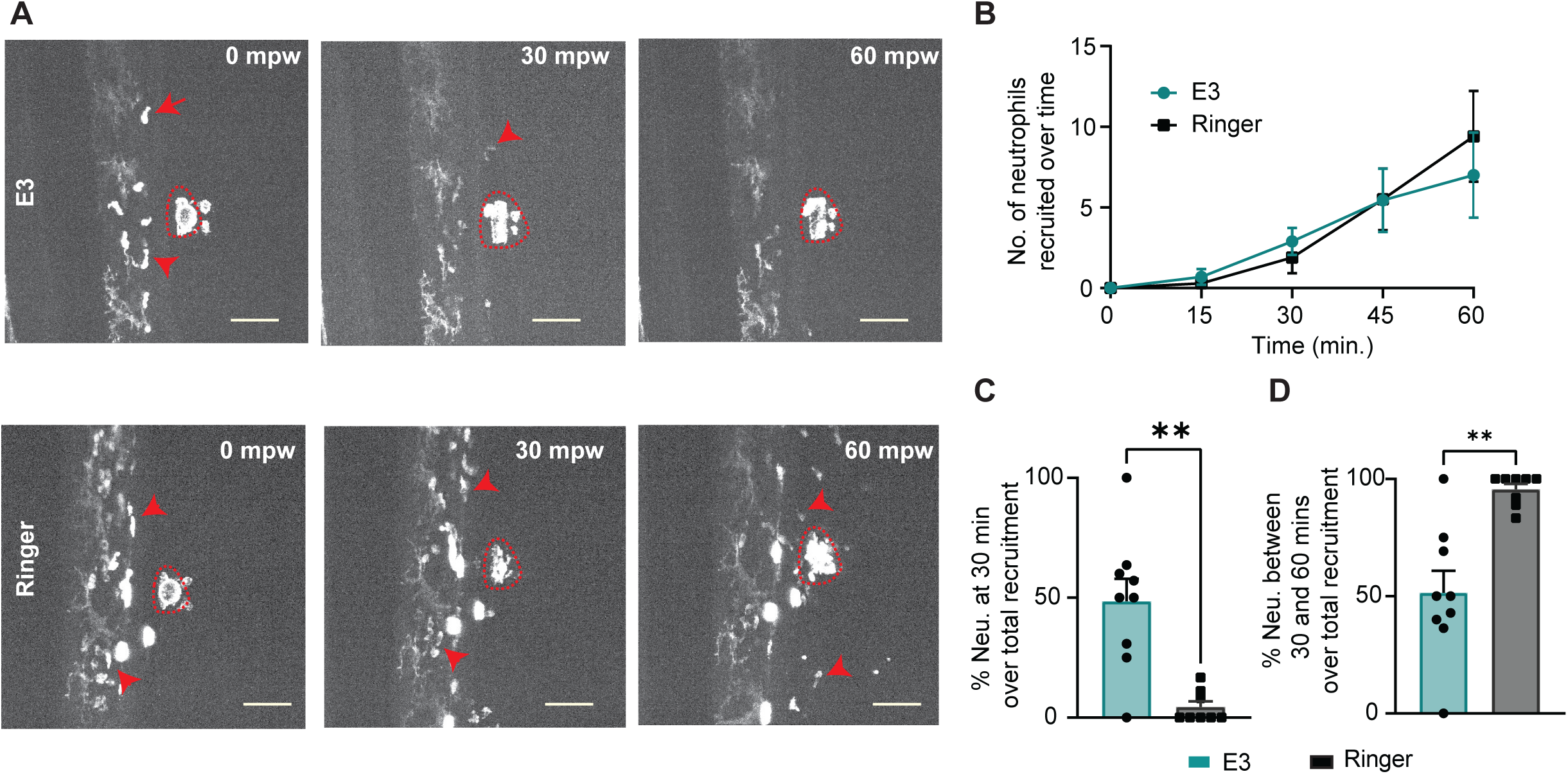
Pseudomonas infection in isotonic solutions elicits delayed neutrophil recruitment at wound-infection sites. A. Time lapse images of representative movies of zebrafish larvae at the indicated time points relative to the time of LW and PAO1 infection in hypotonic (E3) and isotonic (Ringer) buffers. 0 mpw *(left panel)*, 30 mpw *(middle panel),* and 60 mpw *(right panel)*. Red arrows indicate neutrophils and red dotted line depict the site of wound-infection. LW: laser wound, mpw: minutes post wounding. Scale bar 50 μm. *For representative images at 60mpw, the actual time would be 59 mpw*. B. Number of neutrophils recruited at the wound-infection site in zebrafish larvae incubated in E3 and Ringer buffer containing PAO1 after laser wound (LW). n=10 (E3), n=10 (Ringer). Two-way ANOVA with Tukey’s multiple comparisons test. No statistical differences in neutrophil numbers in comparisons across the three buffers. Within the same buffer, there was a difference between the number of neutrophils recruited at 60 min. over 15 min. C. Percentage of neutrophils recruited at the wound-infection site in the early phase (30 minutes) over the cumulative neutrophils recruited after 60mpw. n=9 (E3), n=8 (Ringer). Unpaired t test with Welch’s correction. **P < 0.01. D. Percentage of neutrophils recruited at the wound-infection site in the late phase (between 31 min-60 min) over the cumulative neutrophils recruited after 60mpw. n=9 (E3), n=9 (Ringer). Unpaired t test with Welch’s correction. **P < 0.01.

## Discussion

Here we show that attenuation of osmotic differential detection across the epithelial barrier increases susceptibility to *Pseudomonas aeruginosa* infection. Exposure of wounds to isotonic conditions resulted in enhanced bacterial colonization as early as 30 minutes post infection. Changes in bacterial fitness and virulence within this time scale did not account for the enhanced susceptibility. Rather, a delay in wound closure in isotonic conditions and sub-optimal neutrophil recruitment are likely factors for increased microbial colonisation and vulnerability of the animals. Our results establish the relevance of earlier findings using *P. aeruginosa* microinjection in the context of natural wound infection [21]. Our findings also inform the design of experimental wound infection models in zebrafish, allowing further investigation of wound infection mechanisms in this tractable system.

It remains unclear whether osmotic surveillance influences all wound infections in a similar way. Previous studies reported that damage detection through osmotic surveillance compromised *Candida albicans* wound infection outcome [28]. Mechanistically, this was attributed to keratinocyte movement in response to hypotonic medium detection, which, rather than limiting microbial colonisation, appeared to favour tissue spreading of *C. albicans*. These discrepancies suggest that the role of osmotic surveillance in wound infection outcome depends on the type of invading microbe. For example, *P. aeruginosa* bacteria are highly mobile [29] and thus may rely less on host tissue movements for spreading than *C. albicans*.

*Pseudomonas aeruginosa* is frequently isolated from hospital water systems, sinks, taps, and moist environments [30, 31], and is a leading cause of wound infections. The role of osmotic surveillance in determining susceptibility to infections is relevant in the study of waterborne and hospital-acquired wound infections because barrier disruption not only allows microbial entry but also alters the osmotic environment that initiates wound-sealing and immune responses. This could be an important component for consideration in designing wound management practices. Notably, a previous study found no difference in the infection rate of soft-tissue wounds irrigated with hypotonic tap water or sterile saline solutions but exhibited a clinical trend towards fewer wound infections in the tap water group [32]. The efficacy of the use of hypochlorous acid (HOCl), a hypotonic solution, in the accelerated healing of burn wounds compared to saline solutions [33] indicates that osmotic surveillance mechanisms could be important in wound healing. However, as with the differences in the effect of hypotonic exposure to pathogen growth between *Pseudomonas* and *Candida*, the use of hypotonic solutions in wound management would be context dependent. In the treatment of diabetic foot ulcers, for example, hypertonic saline solution exposure resulted in improved wound healing [34]. Thus, differential wound management may be required based on microbial species, virulence determinants, and host determinants.

Together, our findings demonstrate that the external liquid environment of the wound is a co-factor for wound susceptibility by *Pseudomonas aeruginosa*, alongside host and microbe determinants.

## Acknowledgements

The *P. aeruginosa* strain was kindly provided by Prof. Martin Welch, Department of Biochemistry, University of Cambridge I.W. and S.M.B. were supported by the European Research Council (ERC) under the Horizon 2020 program and UKRI, Grant agreement No. EP/Y02799X/1 awarded to M.S. H.A.W. was supported by an MRC DTP studentship (MR/N013433/1) and a Leverhulme Trust grant (RPG-2021-226). A.D. was supported by a MIRES scholarship from IdEx Université Paris Cité ANR-18-IDEX-0001, funded by the French Government through its ’Investments for the Future’ program. The authors gratefully acknowledge the help by the Aquatics facility at the Department of Physiology, Development and Neuroscience, and the Microscopy Bioscience Platform, University of Cambridge for their support and assistance in this work.

## Materials and Methods

### Experimental model and subject details

All zebrafish were maintained in accordance with UK Home Office regulations under the UK Animals (Scientific Procedures) Act 1986 (PPL number PP4458156), with approval from the University Biomedical Service Committee. Adult fish were maintained and bred following established protocols^69^. Briefly, zebrafish were bred and maintained under standard conditions at 28.5 ± 0.5 °C on a 14 h light:10 h dark cycle. Embryos were collected from natural spawnings at 3 hours post-fertilization (hpf), bleached for 5 minutes in 0.003% sodium hypochlorite (NaOCl; Cleanline, CL3013), and rinsed three times with E3 medium. They were subsequently incubated at 28 °C in E3 medium supplemented with 0.1 µg /ml methylene blue (Sigma-Aldrich, M9140-25G). For imaging experiments, E3 medium was additionally supplemented with 0.003% 1-phenyl-2-thiourea (PTU; Sigma Aldrich, P7629) to inhibit pigmentation and preserve optical transparency.

The following transgenic zebrafish lines were used: Tg(*lyz*:GCamp6F)*^cu104^*

### Bacterial culture and preparation

For *P. aeruginosa* infection, PAO1 strain was used^9^. One day prior to infection, a single colony of *P. aeruginosa* was obtained from a four-way streak on Pseudomonas Isolation Agar (PIA; BD Difco, 292710) supplemented with cetrimide and nalidixic acid (E&O Laboratories Ltd, LS0006) and glycerol. The colony was inoculated into 5 ml of antibiotic-free LB broth and cultured at 37 °C with shaking for 24 h. On the day of the experiment, the overnight culture was diluted in fresh LB and incubated to mid-logarithmic phase (OD₆₀₀ = 0.6–0.8). Bacterial density was adjusted to 3 × 10⁵ CFU/mL using spectrophotometric estimation. Bacteria were pelleted at 1700 × *g* for five minutes and washed twice with PBS to remove residual growth medium, then resuspended in PBS.

### Tail transection, wound infection and survival assay

Zebrafish larvae were transferred to a Petri dish containing Ringer’s solution and 160-200 mg/L MS-222. Tail transection was done using a surgical blade (Swann Morton, 0510) at the distal boundary of the notochord, avoiding injury to the notochord. Non-transected larvae served as controls. Immediately after tail transection, larvae were transferred to a six-well plate containing specific buffers (E3, isotonic E3, Ringer’s solution) with 3 × 10⁵ CFU/mL *P. aeruginosa* and incubated at 32 °C for 3 h. After incubation, larvae were washed five times in E3 medium (without methylene blue). Individual larvae were then placed in wells of a 24-well plate containing 1 mL E3 (without methylene blue) and monitored for survival and disease progression. The larvae were kept at 32 °C over the 24 h observation period.

Survival was assessed immediately after washing, at 3 hpi, and 24 hpi. Assessment was also done at intermediate time points (7 hpi and 10 hpi) when experiment was done for 24 h. Phenotypic disease scoring was based on behavioural responses, transparency, tissue integrity, circulation, and survival. Between observation timepoints, larvae were returned to 32 °C. A score of 4 was defined as death of the organism that was determined by the absence of heartbeat as observed under the microscope.

### Bacterial burden quantification

At indicated timepoints, zebrafish larvae from each condition were pooled into groups of three larvae and homogenised in 300 μL PBS containing 0.1% v/v Triton X-100 (VWR, 28817.295) using sterile pestles. Serial dilutions (up to 10⁻³) were prepared in PBS and plated on PIA supplemented with cetrimide and nalidixic acid (as per manufacturer’s instructions) for *P. aeruginosa.* CFU were then counted after 24 hours incubation at 37°C.

### Effect of buffers on bacteria

Equal number of mid-log phase bacteria (PAO1) based on spectrophotometric estimation was incubated in E3, isotonic E3 and Ringer solution for 30 minutes at 32°C (the temperature at which zebrafish larval infections are performed). Serial dilutions were made in PBS and plated on PIA supplemented with cetrimide and nalidixic acid (as per manufacturer’s instructions) for *P. aeruginosa.* CFU were then counted after 24 hours incubation at 37°C.

### Two-photon imaging and laser wounding

Two photon imaging was performed in Tg(lyz:GCaMP6f)^cu104^ larvae. Transgenic larvae expressing the calcium indicator GCaMP6f were screened to ensure high reporter expression. Larvae exhibiting strong fluorescence and normal morphology were selected for imaging and infection. Selected larvae were anesthetized and mounted in a mixture of 1.2% low-melting-point agarose (Invitrogen, 16520) and the medium (E3, isotonic E3, Ringer) containing 160-200 mg/L MS-222. Imaging under sterile conditions was performed as previously described^54^. For sterile imaging, larvae were mounted on a 35 mm glass-bottom dish with a 14 mm microwell #1.5 cover glass. Excess agarose was removed before solidification to embed the larvae in a thin layer. The mounted larva was bathed in specific buffer as required. To image larvae upon laser wound and infection, a customised imaging chamber was used consists of a metal mould with grooves for cover slips on both sides. Adding both the cover slips created a chamber wherein the mounting was done as described previously. The mounted larva was bathed in specific buffer with required bacteria concentrations.

Laser ablation and time-lapse imaging were performed on a LaVision TriM Scope multiphoton microscope equipped with an electro-optic modulator for rapid power modulation. A Spectra-Physics Insight DeepSee dual-line laser was tuned to 900 nm for imaging and 1,040 nm for ablation, with imaging power set to ∼500 mW at the specimen plane. Image acquisition was controlled using ImSpector Pro software (version 5.0.284.0; LaVision Biotec, ©1998–2016). Two-photon imaging was conducted with a 25×/NA 1.05 water-dipping objective. A 40-μm-diameter region of interest was defined on a single superficial focal plane (240 nm/pixel, 15 μs dwell time). Z-stacks (∼20 planes, 2 μm step size) were acquired every 20 s. Laser wounding was initiated after a 2 min pre-wound baseline, followed by 1 h of post-wound imaging. Focus was maintained throughout acquisition. Image stacks were processed in Fiji using maximum-intensity z-projections to generate neutrophil-swarming time-lapse movies for subsequent analysis.

### Image analysis

#### Cumulative number of neutrophils arriving at the wound

The cumulative number of neutrophils that reach the wound margin were calculated manually by counting the fluorescent neutrophils reaching the wound in blinded datasets using ImageJ/Fiji.

#### Measuring rate of wound closure

Time-lapse movies of laser wounding in zebrafish larvae (3dpf) in three buffers (E3, E3 isotonic and Ringer) were acquired as mentioned above. All movies were blinded before analysis of wound closure. Each time lapse movie was opened using ImageJ/Fiji. The first slice in which the impact of the wound is visible is noted. For analysis of the wound closure, the slice after the first visible impact of the wound is selected as the time of wound (t=0 mins). Using the freehand selection tool, the margin of the wound is selected as the region of interest (ROI) and the area of the ROI measured. Similarly, the wound region is outlined in all subsequent time points, and their corresponding area of ROI is measured (until t=20 mins). Exporting the area onto an excel sheet, the area at each time point is normalized by the initial area (t=0). The resulting value is plotted over time. A non-linear regression (curve fit) analysis was done using GraphPad Prism. Briefly, the one phase decay model with the least squares regression method was employed. The comparison was done to check if the best-fit values for the three conditions differ for all parameters (Y0, plateau and K).

### Statistical analyses

All statistical tests were performed in Prism10 (GraphPad Software, La Jolla, CA). The statistical test and the n number are indicated in the figure legends. The error bars show standard error of the mean except for bacterial burden data where error bars represent 95% confidence intervals of the median. Where the distribution was verified as normal, outliers were removed by applying Rout test. Live imaging experiments were acquired in minimum three independent experiments. A normal distribution test was performed prior to any *t*-test statistical analyses. Normal distribution was assessed using *Shapiro-Wilk test*.

