## Supplementary Material for "Tissue osmotic surveillance shapes susceptibility of zebrafish to wound infections"

### Supplementary Movie Legends

**Movie S1 Examples of wound closure and neutrophil recruitment to laser wound in E3, E3 isotonic and Ringer medium in sterile conditions**

Examples of neutrophil migration in a Tg(*lyz*:GCamp6F) zebrafish larvae pre and post laser wound in E3 (*left*), E3 isotonic (*middle*) and Ringer (*right*). Frame intervals 20 sec and frame rate is 20 fps. Scale bar 50 µm.

**Movie S2 Examples of neutrophil migration in E3 and Ringer medium containing PAO1**

Examples of wound closure and neutrophil migration in PAO1 infected Tg(*lyz*:GCamp6F) zebrafish larvae post laser wound in E3 and Ringer solutions. Frame intervals 20 sec and frame rate is 20 fps. Scale bar 50 µm.
